# *ATNr*: Allometric trophic models in R

**DOI:** 10.1101/2022.08.26.505404

**Authors:** Benoit Gauzens, Ulrich Brose, Eva Delmas, Emilio Berti

**Author notes:** Benoit Gauzens, Puschstraße 4, 04103 Leipzig.

## Abstract

1. Understanding and predicting how densities of interacting species change over time has been one of the main goals of community ecology, which has become a pressing challenge in the context of global change.
2. We present the R package *ATNr*, which provides an implementation of different versions of Allometric Trophic Network models (Yodzis and Innes (1992)) that simulate the biomass dynamics of trophically interacting species.
3. Relying on *C++* routines, the *ATNr* proposes an efficient and standardized implementation of the different ATNs models.
4. By proposing a set of built in functions ready to use in a language widely used in the community of ecologists, the *ATNr* package offers an easy access to ATN models.

## Introduction

Understanding and predicting how densities of interacting species change over time has been one of the main goals of community ecology and that has become a pressing challenge in the context of global change, which asks for predictions of how anthropogenic and natural changes in environmental conditions will influence the functioning of future communities. Indeed, species biomass density influences ecological processes (e.g. energy fluxes), species persistence, and ecosystem stability (Kramer et al. (2018); Gauzens et al. (2019)). Therefore, understanding how species’ biomass changes over time is necessary for basic ecological research as well as applied management and conservation. However, tackling this issue with experimental approaches faces several limitations, as collecting temporal datasets on the dynamics of species from complex communities over long period of times is nearly impossible.

Due to such empirical limitations, predictions on dynamic of species densities are usually achieved using theoretical models that implement ordinary differential equations (ODEs) describing the rate of change in species’ densities depending on their interactions with the other species in the community (Volterra, 1927; Sauve and Barraquand, 2020; Boit et al., 2012). Parametrising such models, however, can easily become unmanageable for large communities, due to the required quantification of species’ biological rates and inter-specific interaction strengths. To solve this problem, Yodzis & Innes (1992) proposed a “bio-energetic” model that quantifies the key parameters driving species dynamics using allometric relationships with body mass (Williams et al., 2007). This greatly simplified the problem of model parametrisation, allowing to use body mass in order to derive the other parameters.

The general approach of the bio-energetic model, extended by the development of metabolic theory (Brown et al., 2004), led to a new family of models, called Allometric Trophic Network models (ATNs, Martinez 2020), as their parametrisation heavily relies on the body mass of species. In practice, allometric relationship are used to relate the body mass of species to several of the biological processes driving feeding rates and species dynamics (for instance, attack rate, handling time, and metabolic rates are all function of species’ body mass) and, in some cases, to estimate the presence or absence of trophic interactions Schneider et al. (2016). Interestingly, ATNs have been shown to have the potential to represent ecosystem complexity realistically, predicting accurately community dynamics (Boit et al., 2012; Curtsdotter et al., 2019) and responses to environmental gradients (Fussmann et al., 2014). ATN models provide thus a practical compromise between biological complexity (their parametrisation is eased by the use of few easily measurable traits, i.e. species’ body mass) and realism in order to model density dynamics of interacting species.

Three major ATN models are currently used today, hereafter referred as *unscaled, scaled*, and *unscaled with nutrients*. These models differ in the possibility of including nutrient dynamics or in the way the different biological rates underlying species growth rate are determined: biological rates can be either estimated from empirical observation without further transformations or scaled to a desired level for the sake of comparability. More formally, the *unscaled* model is based on untransformed biological rates (such as attack rate and handling time) derived using allometries. The *unscaled with nutrients* model relies on the same set of assumptions as the *scaled* version, but it explicitly includes the dynamics of nutrients and their interactions with plant species. The *scaled* model, on the other hand, scales the biological rates to the growth rate of the smallest basal species). Together, these models permit to simulate species’ density dynamics in a variety of cases, e.g. when nutrients are explicitly accounted for and when the interest is in the untransformed biological rates or in their relative quantities. Currently, however, the application of ATN models is hindered by a lack of a standard framework, with studies using different implementations for the same underlying mathematical model.

An important step towards a common framework was recently achieved by Delmas et al. (2017), who developed a standard implementation for ATNs in the Julia programming language. Yet, this implementation focuses on a specific class of ATNs (*scaled*) and in a programming language that is unfamiliar to many ecologists. Here, we present the new R package *ATNr*, which provides a standard implementation for the three major classes of ATN models. Importantly, in *ATNr*, the core of the calculations are evaluated by *C++* routines called from within the R environment, providing a standard framework that is familiar to many ecologists while having a high computational performance.

### 1 ATN models

All ATN models available in *ATNr* describe the biomass dynamic of species by estimating their net growth rates over time. *ATNr* relies on two general assumptions: i) biomass production of species are positively affected by what they consume, i.e. the models are based on energetic transfers between resources and their consumers (plant species can either consume nutrients or have intrinsic logistic growth rates); ii) loss rates of species are driven by their consumers and by metabolic costs. In this section we cover the main aspects of the three ATN models contained in *ATNr*, highlighting the major differences between them. More details and the units of the different variables can be found in the package vignette (`vignette(`“`model_descriptions`”`, package =` “`ATNr`”)).

#### 1.1 Species growth rate

Growth rate of species *i* is defined as the change of its biomass (*B*_*i*_) over time: 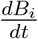. For basal species, the growth rate depends on their net intrinsic growth rate minus their metabolic losses and the negative effects of their consumers:

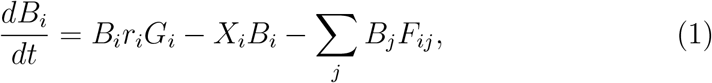

where *r*_*i*_ is the maximum, mass-specific growth rate, *G*_*i*_ defines the proportion of *r*_*i*_ that is realised such as *B*_*i*_*r*_*i*_*G*_*i*_ corresponds the net growth rate. *X*_*i*_ is the per-biomass metabolic losses, and *F*_*ij*_ is the per-biomass feeding rate of consumer *j* for resource *i* (*F*_*ij*_ = 0 if *j* does not feed on *i*).

For non-basal species, net intrinsic growth rate is substituted in eq. 1 by the biomass gained from resources multiplied by the assimilation efficiency of the resource *j* (*e*_*j*_):

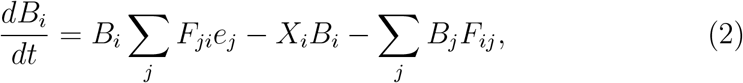

These equations define allometric models as the terms involved are parametrized using scaling relationships with body mass of species (*M*). For instance, *r*_*i*_ and *X*_*i*_ both follow a power law that is a function of the body mass *M*_*i*_ of species *i*:

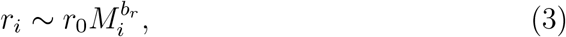

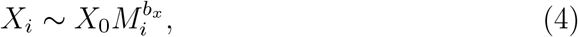

where *r*_0_ and *X*_0_ are normalisation constants and *b*_*r*_ and *b*_*x*_ the scaling exponents.

The three different versions of ATN models proposed in the package all derive from this set of equations, and only differ in how the feeding rates *F*_*ij*_ (also called functional response) are specified and how the net growth rates of basal species are calculated. For instance, a major difference in the calculation of *G*_*i*_ among models is whether or not the dynamic of the nutrient pool is considered.

Here we present the formulation of feeding rate and net growth rate used by the different models and depict how the different variables are usually defined by proposing a default parametrisation based on what is used in the literature. As a convention, for all parameters that depend on both resources and consumers, like *F*_*ij*_, the first index refers to the resource and the second to the consumer. We provide the default parametrisations of model terms commonly used in literature in the package vignette.

#### 1.2 Unscaled ATN

This ATN implements the model as in Binzer et al. (2016). In *ATNr* it is called *Unscaled* because the biological rates are used in their untransformed form (contrary to the *Scaled* ATN, see below). The functional response and growth rate are defined as:

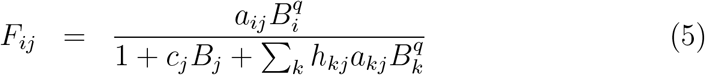

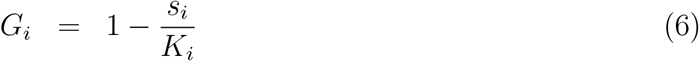

where *a*_*ij*_ is the attack rate of *j* on *i, q* is the Hill exponent determining the shape of the functional response (for type II, *q* = 1; for type III, *q* ∈]1, 2]), *c*_*j*_ defines the intra-specific interference competition, *h*_*ij*_ is the handling time of *j* on *i, K*_*i*_ is basal species *i* carrying capacity, and *s*_*i*_ is a parameter defining the ratio between inter- and intra-specific competitions for resources among basal species.

#### 1.3 Scaled ATN

This ATN implements the model as in Delmas et al. (2017). It is called *Scaled* because the biological rates of species are scaled to the growth rate of the smallest basal species. The consequence of this scaling is that time is now defined as the inverse of the growth rate of the smallest producer species and the outcome of the model is to be interpreted on a relative time scale. While this may seem unnecessary complicated, it allows to work at a scale adapted to the study system and to directly compare processes that happen on very different time scales (e.g. phytoplankton bloom and forest growth; Williams et al. 2007). In particular, *r*_*i*_ and *x*_*i*_ from eq. 1 and 2 are divided by the intrinsic growth rate and metabolic cost of the basal species with the smallest body mass. When species are sorted by body mass, the first one is the smallest species and: 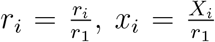. Then, it introduces the maximum feeding rate of species relative to their metabolic rate *y*_*i*_, i.e. the ratio between maximum feeding rate and metabolic rate.

This redefinition of the variables implies a redefinition of the equations describing the dynamics of species:

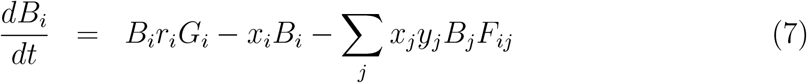

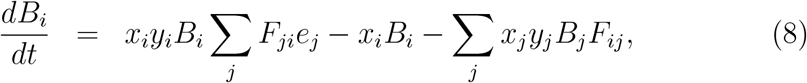

The functional response and growth rate are then defined as:

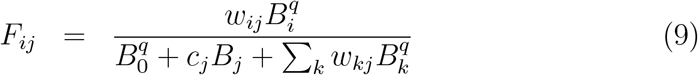

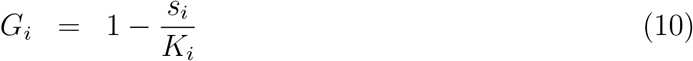

where *w*_*ij*_ is the relative consumption rate of *j* on *i*, such that ∑_*i*_ *w*_*ij*_ = 1.

#### 1.4 Unscaled ATN with nutrients

This ATN implements the model as in Schneider et al. (2016). It differs from the *Unscaled* in how the attack rate is defined and, more importantly, by allowing growth rate of plants to be determined by a nutrient pool which dynamic is explicitly modeled. The functional response and growth rate are defined as:

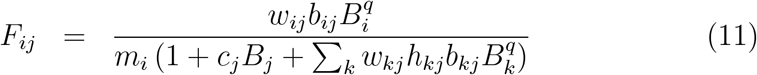

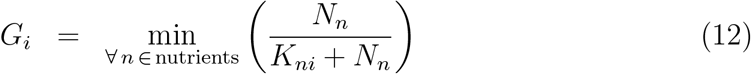

where *b*_*ij*_ is the resource-specific capture coefficient, *m*_*i*_ is body mass of *i, N*_*n*_ is the concentration of nutrient *n*, and *K*_*ni*_ is the nutrient uptake efficiency of basal species *i* on nutrient *n*, smaller for higher efficiencies. The change of nutrient concentration over time is described by another set of differential equations:

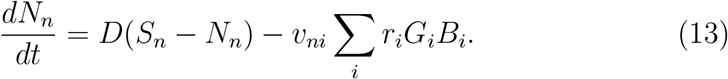

Here, *D* is the global turnover rate that determines the rate by which the nutrients are refreshed, *S*_*n*_ is the maximal concentration of nutrient *n*, and *v*_*ni*_ sets the relative content of nutrient *n* in plant *i*.

### 2 Using the ATNr package

The *ATNr* package consists of compiled *C++* routines, which define the ATN models, together with R functions to interface with the instances of the ATNs and utility functions to solve the ordinary differential equations (ODE) and plot results. The framework of *ATNr* can be divided into three steps (Table 2): 1) create an instance of an ATN model; 2) initialise parameter values (potentially using the default parametrisation proposed); and 3) solve the ODE using an ODE solver.

**Table 1:** Step and *ATNr* functions to create ATN models, specify default parametrisations, and solve the biomass dynamics. *nb*_*s*_ is the number of species, *nb*_*b*_ the number of basal species, *nb*_*n*_ the number of nutrients, *BM* species body mass, *fw* the food web matrix, *model* an instance of ATN model, *temperature* the environmental temperature in Celsius, *L*.*mat* the *L* matrix, *t* the integration time steps, *y* the starting biomasses of species and nutrients, and *x* is the matrix with ODE solutions as returned by `lsoda_wrapper()`.

| Step | ATNr function | required input |
| --- | --- | --- |
| Create an instance of an ATN model | `create_model_Scaled()` | $nb_s, nb_b, BM$ and $fw$ |
| | `create_model_Unscaled()` | $nb_s, nb_b, BM$ and $fw$ |
| | `create_model_Scaled_nuts()` | $nb_s, nb_b, nb_n, BM$ and $fw$ |
| Initialize parameters to their default values | `initialise_default_Scaled()` | $model$ |
| | `initialise_default_Unscaled()` | $model$ and $temperature$ |
| | `initialise_default_Scaled_nuts()` | $model, L.mat$ and $temperature$ |
| Solve ODE and obtain biomass dynamics | `lsoda_wrapper()` | $t, y, model$ |
| Plot biomass dynamics | `plot_odeweb()` | $x, nb_s, model$ |

To create an ATN model, an adjacency matrix of the food web is needed to define consumer-resource interactions together with species body masses to parametrise the ATN. The adjacency matrix can be derived from empirical data or simulated using synthetic food webs. *ATNr* contains two functions to generate synthetic food webs. The first is based on the niche model (Williams and Martinez, 2000; Allesina et al., 2008): `create_niche_model(S, C)`, where *S* is the number of species and *C* the connectance of the network. The second is based on the *L* matrix approach, which uses the consumer-resource body mass ratio (Schneider et al., 2016): `create_Lmatrix(BM, nb_b, Ropt = 100, gamma = 2, th = 0.01)`, where *BM* is a vector of species body masses, *nb_n* the number of basal species, *Ropt* the optimal predator-prey body mass ratio, *γ* the width of the feeding kernel, and *th* is the threshold below which interactions are considered to not be realized.

From species body masses and trophic network, ATN models can be created with the functions `create_model_Scaled()` for *scaled* models, `create_model_Unscaled()` for *unscaled* models, and `create_model_Unscaled_nuts()` for *unscaled with nutrients* models. All three functions generate a *C++* object (formally an instance of the *C++* class) accessible within the *R* environment as an object of class *S4*. Once the model is created, it can be initialised to the default values used in literature (Delmas et al., 2017; Binzer et al., 2016; Schneider et al., 2016) by calling `initialise_default_Scaled()`, `initialise_default_Unscaled()`, and `initialise_default_Unscaled_nuts()`.

Once the ATN models are properly created and initialised, we can solve the ODEs by specifying the integration steps and biomass of species at starting conditions as R vectors and calling an ODE solver. In *ATNr*, the wrapper function `lsoda_wrapper(steps, biomass, atn.model)` calls the lsoda solver from package *deSolve*.

As an example to summarize the general framework of *ATNr*, we compute biomass dynamics for a food web of 20 species generated using the niche model and implementing the *unscaled* ATN model. We start by creating a food web of 20 species using the niche model and specifying a connectance of 0.3:

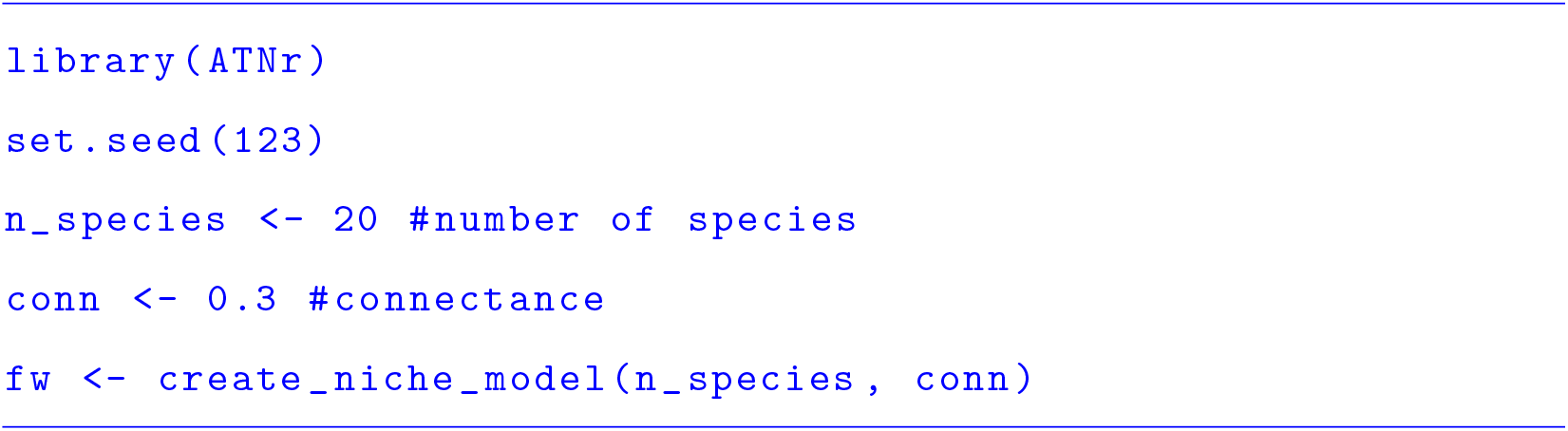

We then calculate the number of basal species and species’ trophic levels, which are used to obtain their body masses:

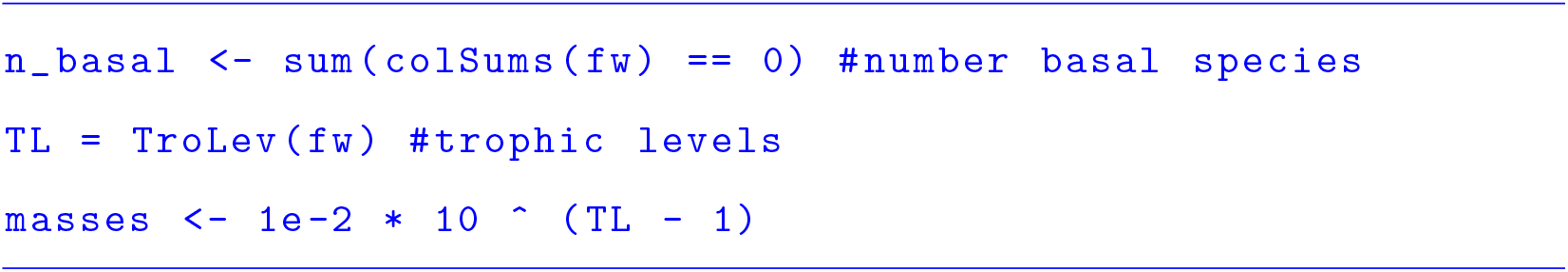

We can now initialise an instance of the *unscaled* ATN model, setting the parameters to their default values:

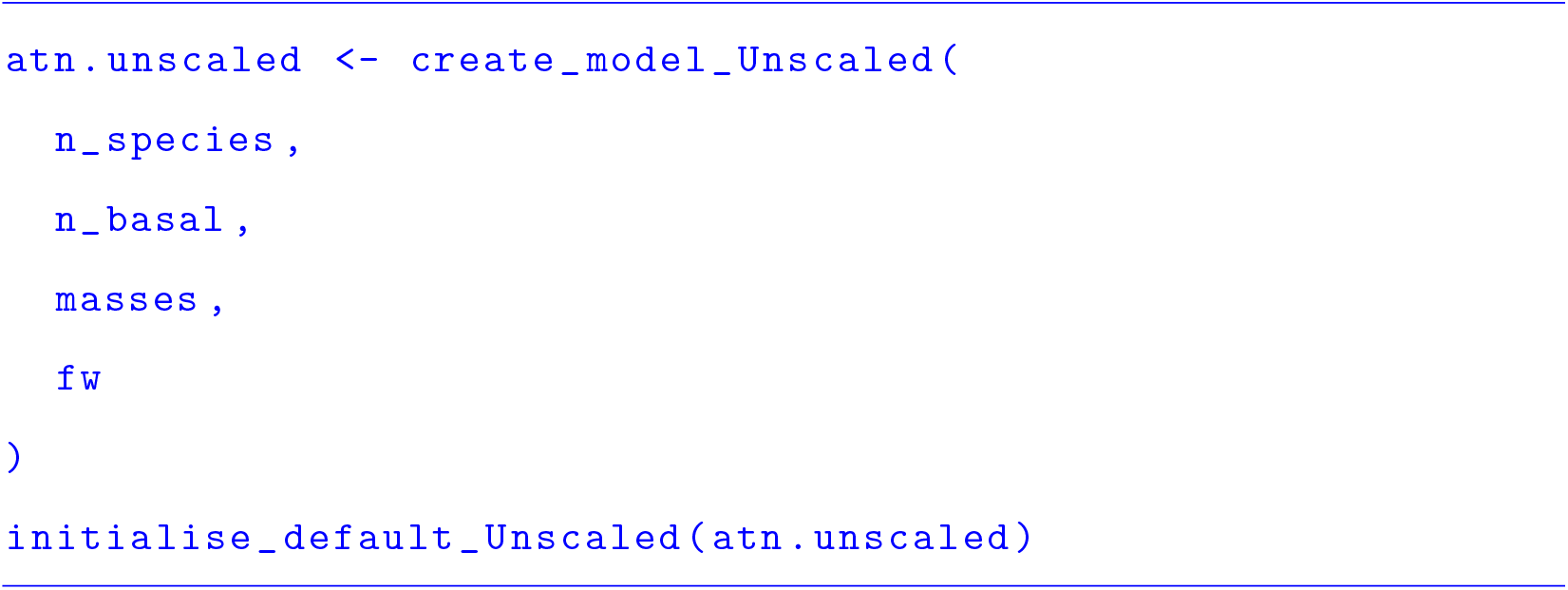

Once we define starting biomasses (here randomly drawn ∈ [2, 3]) and the integration steps, the ODE can be solved:

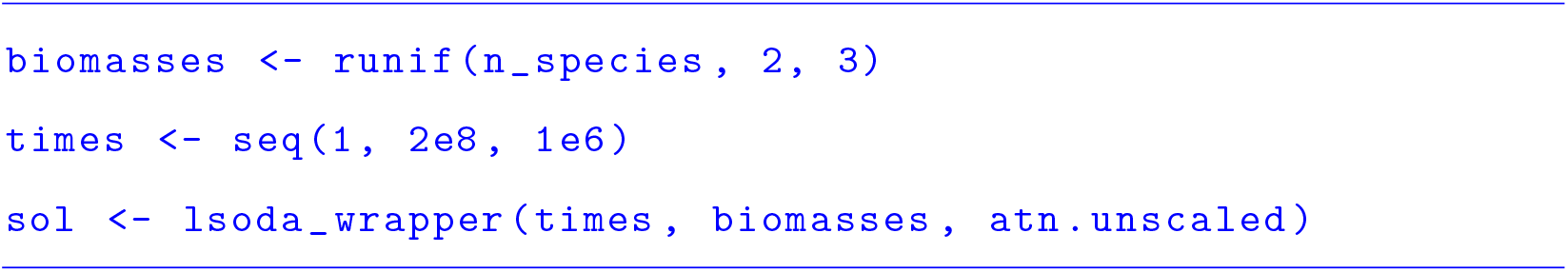

The variable *sol* is a *deSolve* matrix, with *n_species + 1* columns and as many rows as the integration steps. The first column contains the integration steps and the remaining column the biomass of species at each time step. The biomass dynamics can be plotted using *ATNr* function `plot_odeweb()` (Fig. 1):

**Figure 1:**
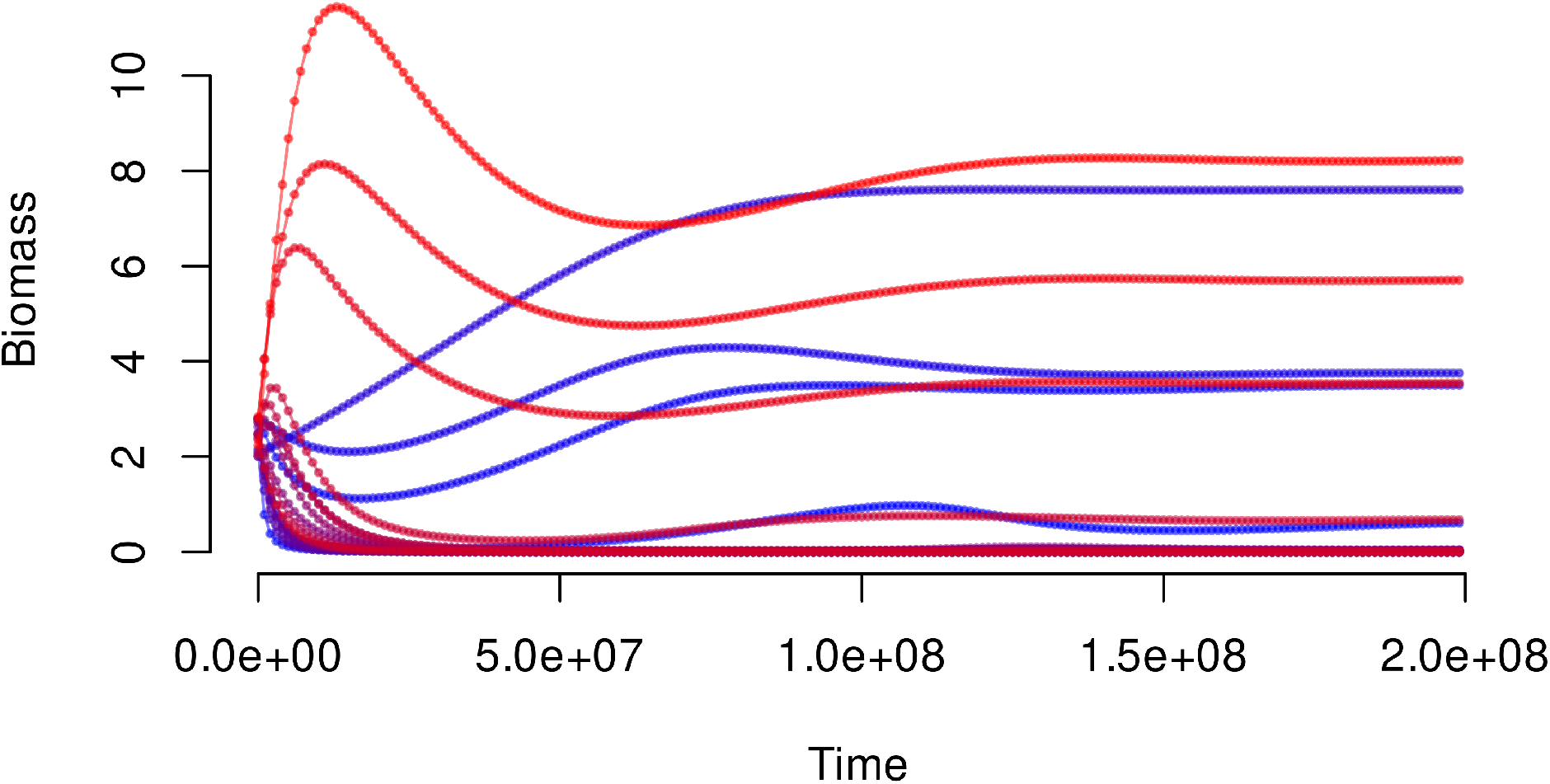
Biomass dynamics as evaluated in the example code. Colors show different species, with dots being the solved dynamics at the pre-defined integration steps.

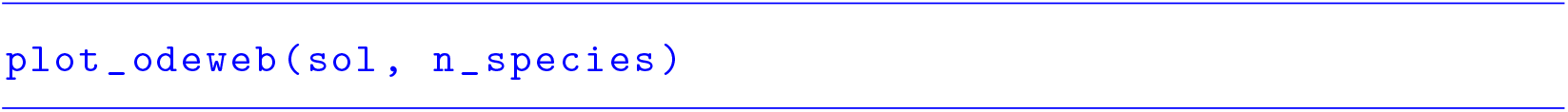

From *sol*, it is straightforward to obtain several metrics with key importance for community ecology, e.g. the number of species that went extinct (i.e. species for which biomass goes below a certain threshold, here 10^−6^):

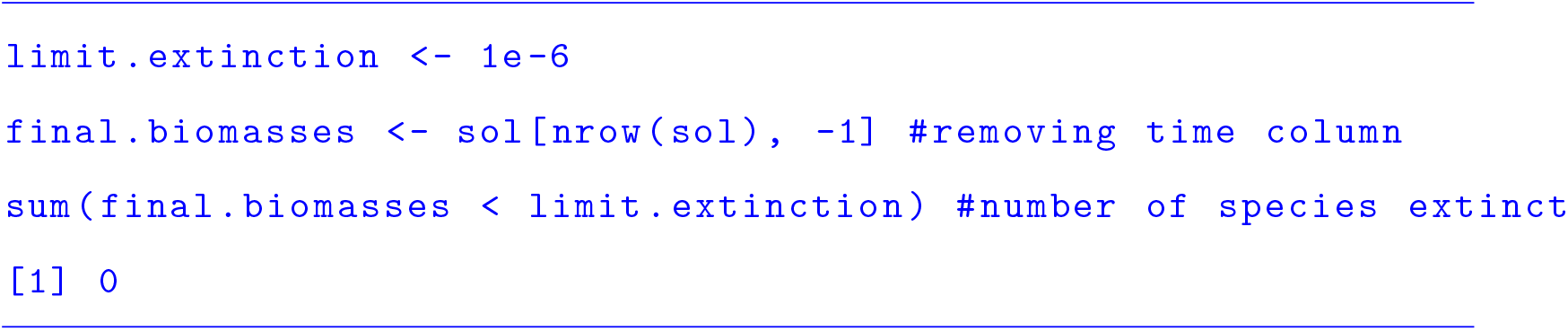

### 3 Applications

To show the potential applications of *ATNr*, we explored two cases: 1) the effects of temperature on species persistence and 2) the paradox of enrichment. These examples are also shown in the package vignette (`vignette(`“`ATNr`”)).

#### 3.1 Effect of temperature on species persistence

*ATNr* makes it relatively easy to vary one parameter to assess its effect on the population dynamics. As example, we focus here on environmental temperature, which has been shown to influence biological rates of organism and to play an important role in their persistence (Gauzens et al., 2020). In *ATNr*, temperature can be specified in the *unscaled* and *unscaled with nutrients* models at initialisation: `initialise_default_Unscaled(model, temperature)` and `initialise_default_Unscaled_nuts(model, L.mat, temperature)`. We first create a food web using the *L* matrix approach:

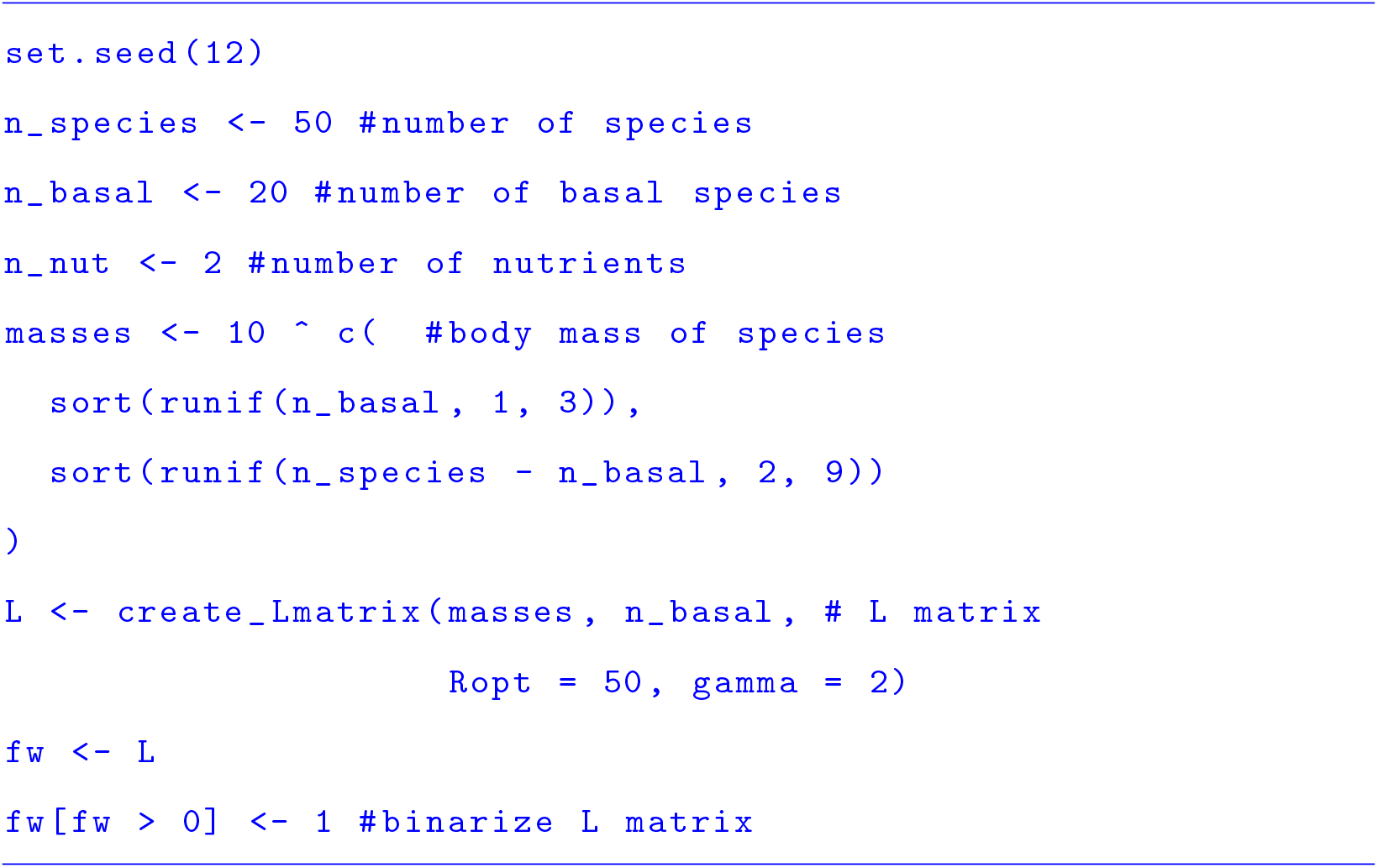

As we are interested in modelling nutrients explicitly, we use here the ATN model including nutrients. We thus create and initialise an instance of the *unscaled with nutrients* ATN model:

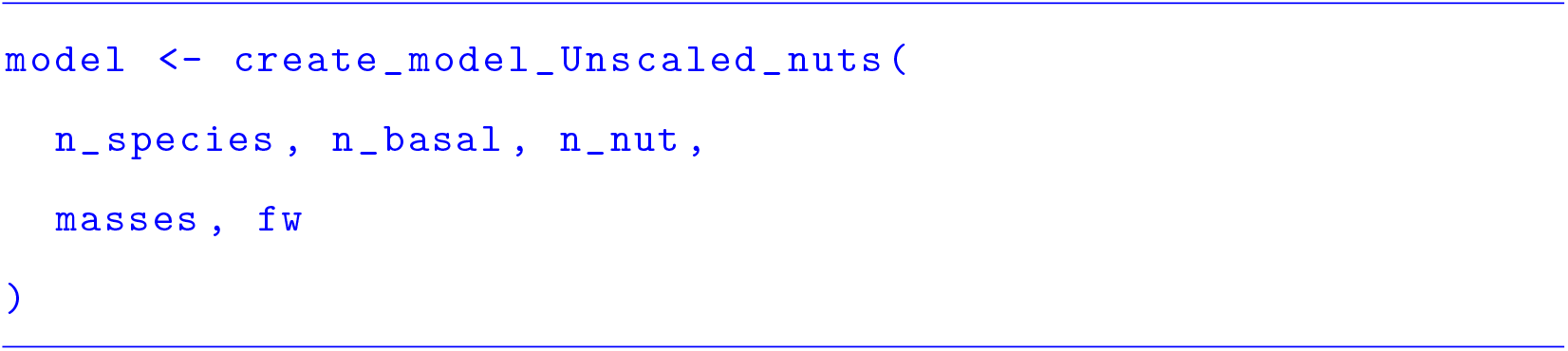

To study the effect of temperature on species persistence, it is sufficient to loop over a temperature range and re-initialise the model instance at every iteration (Fig. 2):

**Figure 2:**
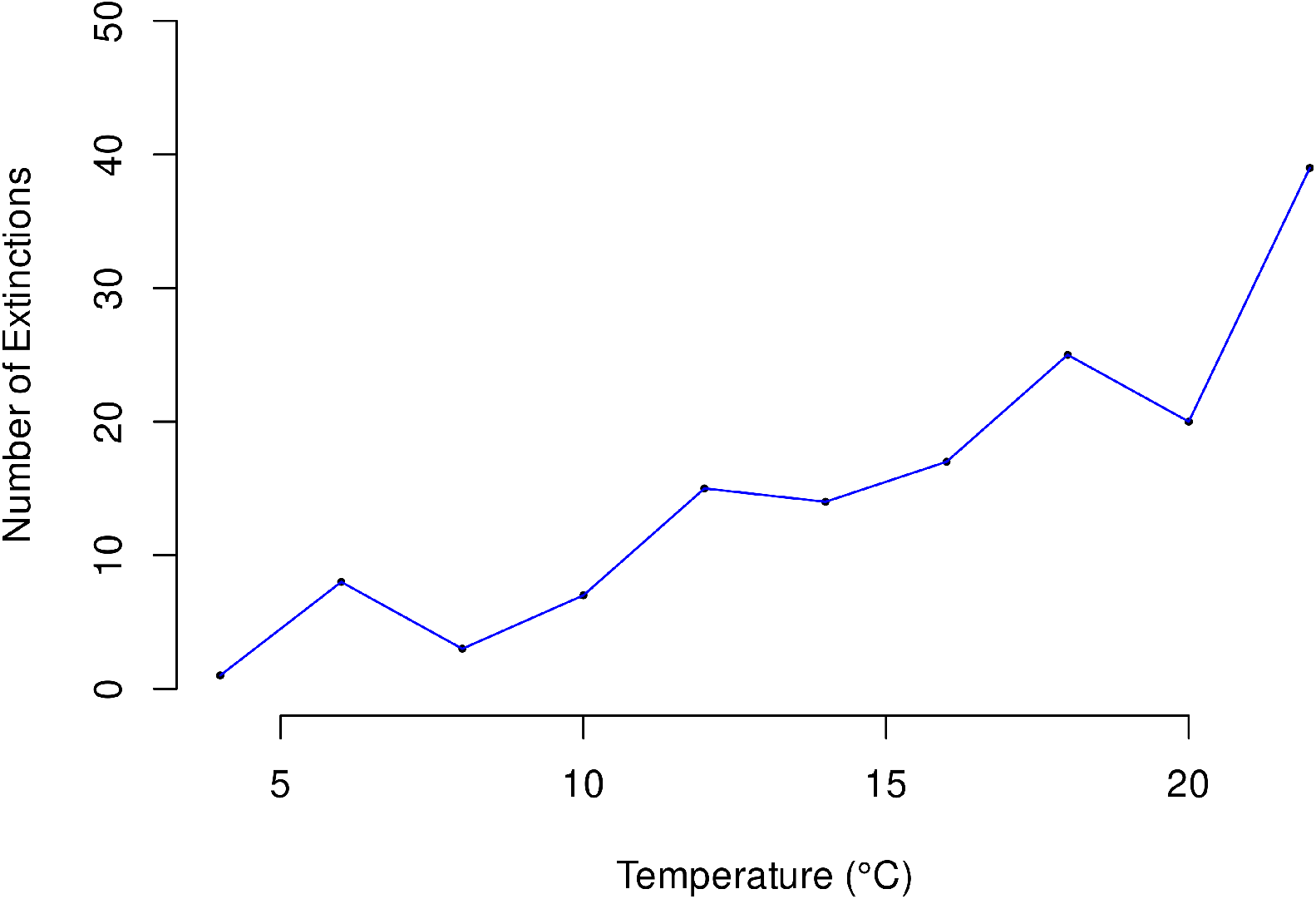
Effect of temperature on species persistence using the *unscaled with nutrients* ATN model. For increasing temperatures, more species go extinct (total species richness = 50).

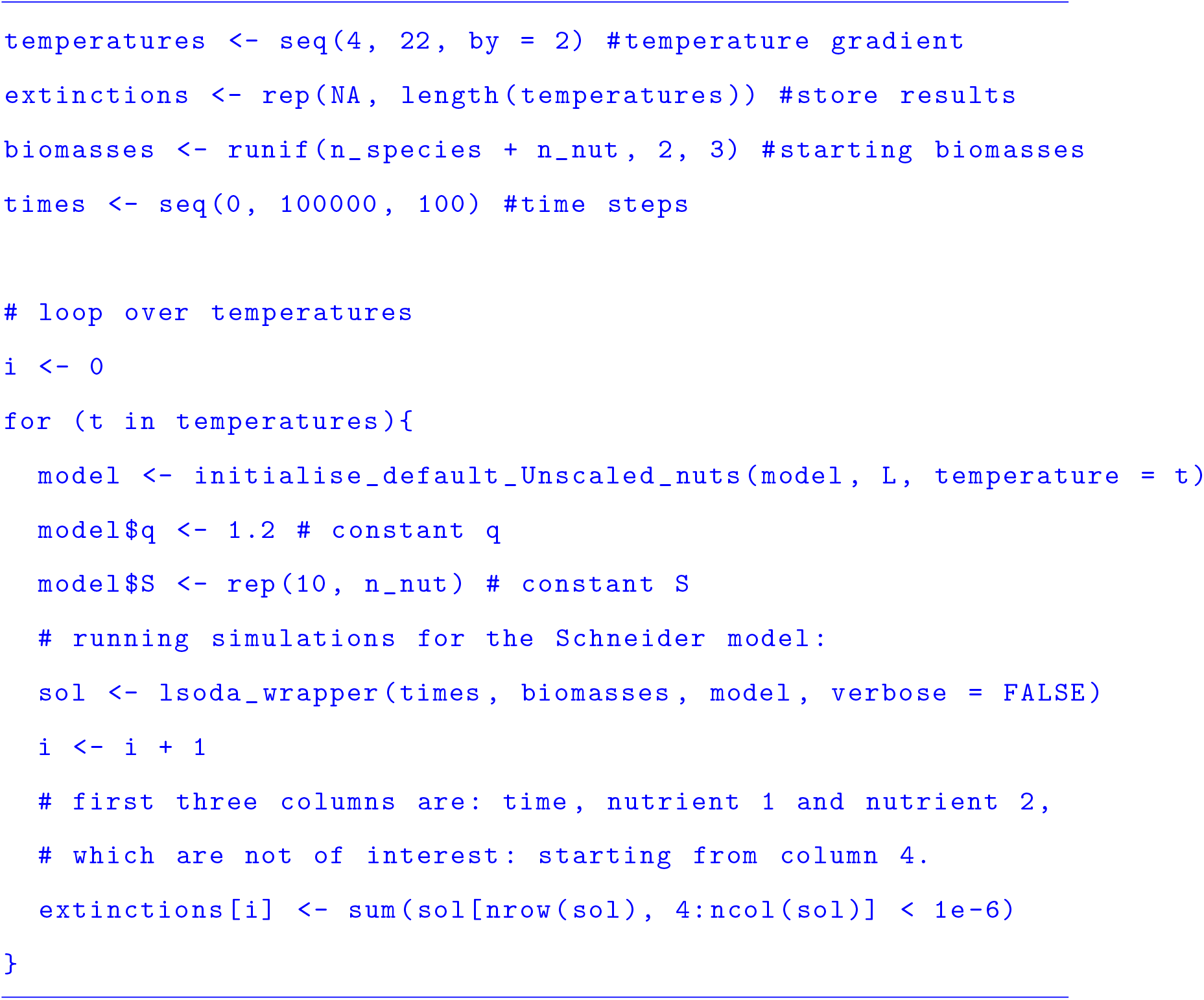

Here, *q* and *S* were fixed to constant values, instead of being randomly sampled, in order to make runs at different temperature comparable.

As a note, we want to point out that in the above example there is no need to create separate instances of the ATN model, as internal parameters (temperature in this case) are changed only through `initialise_default_Unscaled_nuts()`. When this is not the case (e.g. changing number of nutrients or species) a new instance of the ATN model is needed for each simulation.

#### 3.2 Paradox of enrichment

Here, we want to test the paradox of enrichment, i.e. that increasing the carrying capacity of basal species may destabilize population dynamics, by using the *scaled* model. First, we create a food web with 10 species and initialise the model:

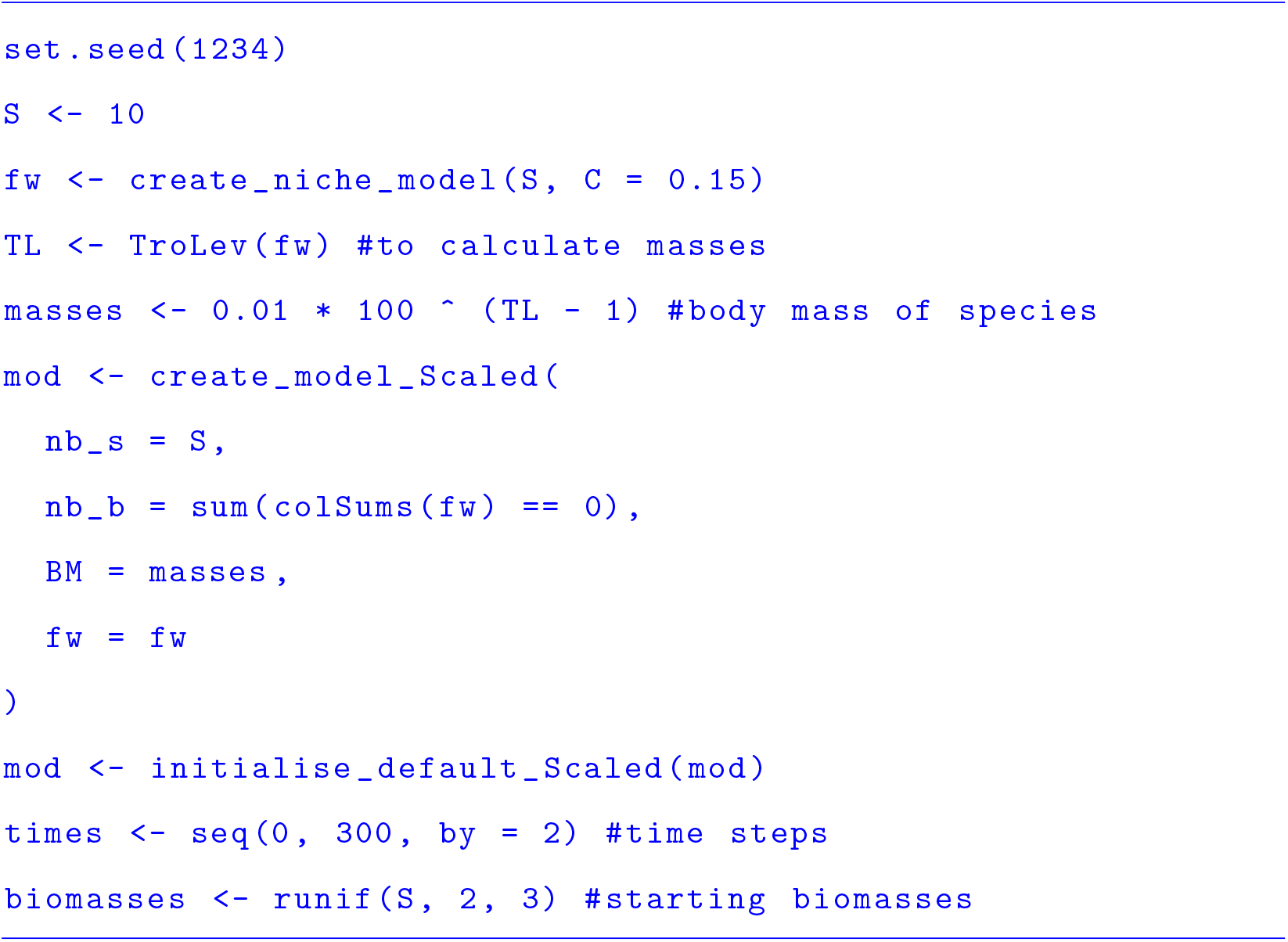

Then, we solve the system specifying the carrying capacity of basal species equal to one (`mod$K <-1`) and then increased this to 10 (`mod$K <-10`; Fig. 3):

**Figure 3:**
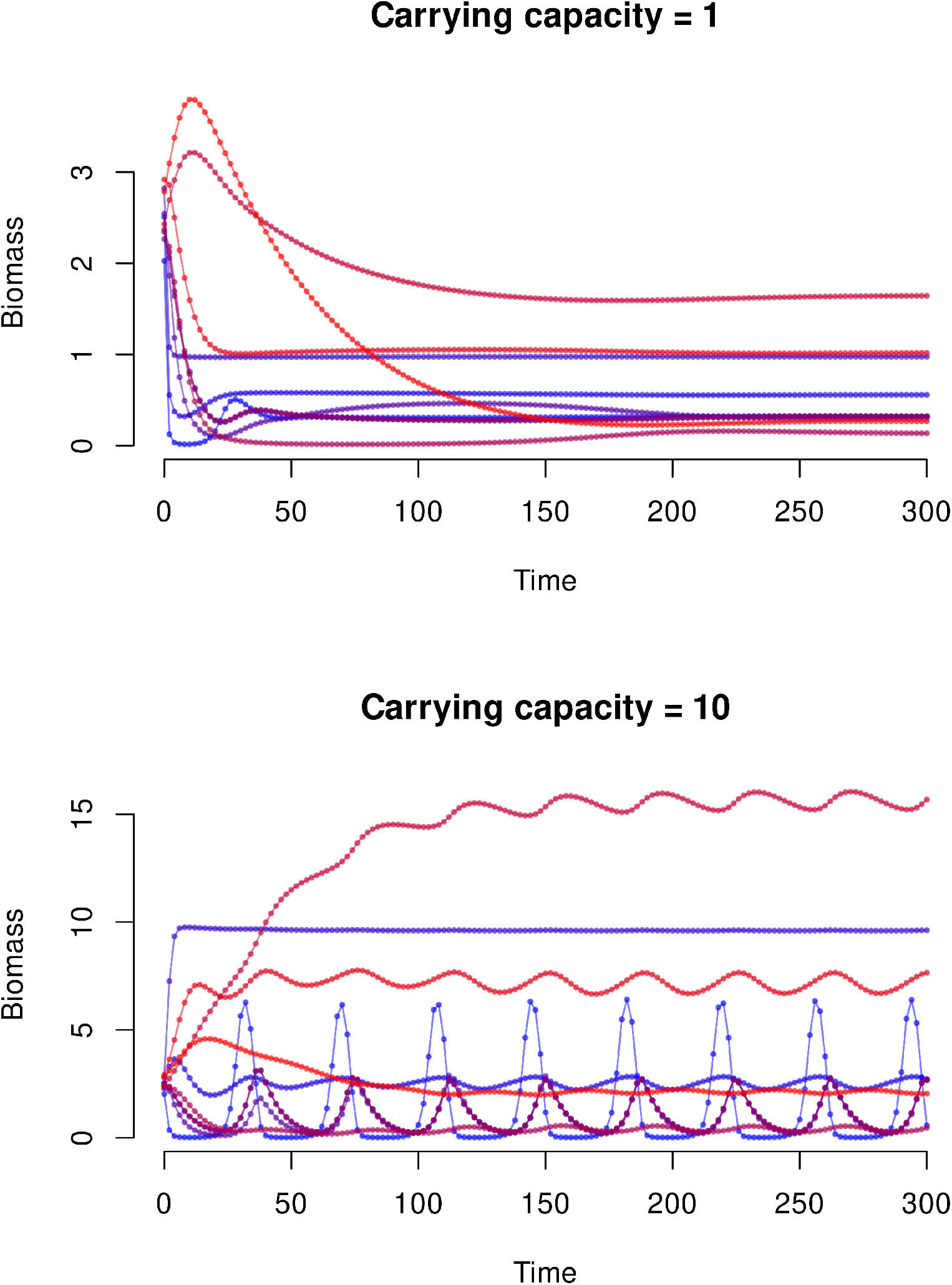
Paradox of enrichment using the *unscaled* ATN model. For higher carrying capacity of basal species (*K = 10*), dynamical oscillations arise, destabilizing the community.

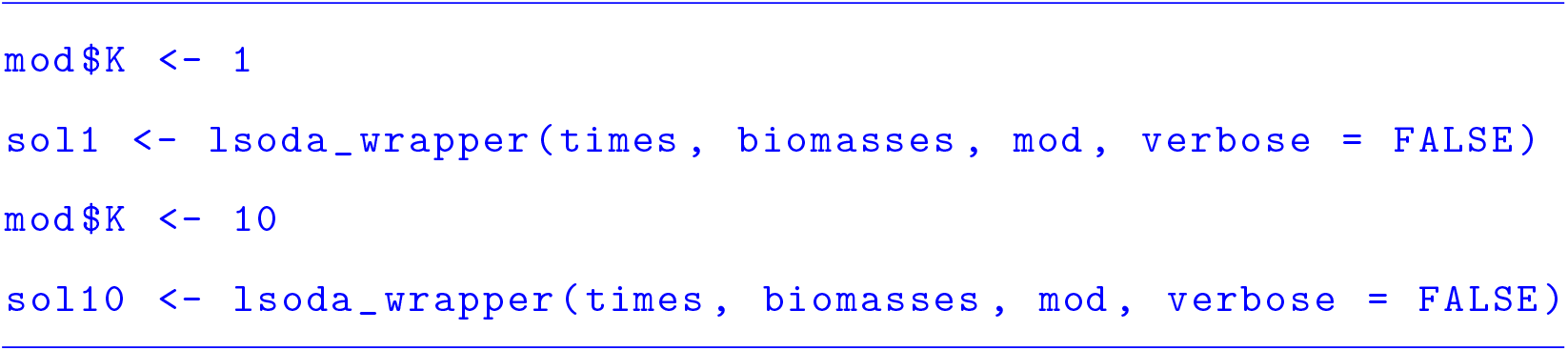

### 4 Conclusion

The R package *ATNr* provides a standardized implementation of different versions of Allometric Trophic Network models, describing the dynamics over time of trophically interacting populations using allometric relationships. *ATNr* provides four major advantages: First, as our implementation relies on *C++* routines, it allows to efficiently use ATN models in order to simulate the dynamics of interacting populations, up to several hundreds of species. Second, *ATNr* unifies the three major ATN models into a single library, providing a standardized implementation that enhances comparisons of different ATNs within a study and compatibility across several projects. Third, it provides an interface to ATN models in the R language, i.e. suited for a variety of users, especially in ecology. Finally, *ATNr* is ready-to-use and makes ATN models available also for novices, while allowing more expert users some degree of flexibility.

## Acknowledgements

BG, EB and UB cknowledge the support of the German Centre for Integrative Biodiversity Research (iDiv) Halle-Jena-Leipzig funded by the German Research Foundation (FZT 118). We thank Daniel Reuman for his helpful comments on the package.

## Author contributions

BG and EB designed the study and developed the package. BG wrote the first draft of the manuscript. All authors made substantial revisions and comments to the manuscript.

## Data accessibility

A beta version of *ATNr* is deposited on CRAN: https://cran.r-project.org/package=ATNr. A development version can be accessed on GitHub: https://github.com/gauzens/ATNr. Should our manuscript be positively evaluated by reviewers, a finalised version will be pushed on CRAN

